# SS-Detect: Development and Validation of a New Strategy for Source-Based Morphometry in Multi-Scanner Studies

**DOI:** 10.1101/2020.09.03.282236

**Authors:** Ruiyang Ge, Shiqing Ding, Tyler Keeling, William G. Honer, Sophia Frangou, Fidel Vila-Rodriguez

**Affiliations:** Non-Invasive Neurostimulation Therapies Laboratory, Department of Psychiatry, University of British Columbia, BC, Canada; Department of Psychiatry, University of British Columbia, BC, Canada; Department of Psychiatry, Icahn School of Medicine at Mount Sinai, NY, USA

**Author notes:** Correspondence to: Ruiyang Ge, PhD, contact address: 2255 Wesbrook Mall, Vancouver, BC V6T 2A1, Canada, OR.

**Keywords:** simulation, source-based morphometry, structural brain pattern, T1-weighted MRI, multi-scanner study

## Abstract

**Background and Purpose:** Source-based morphometry (SBM) has been used in multi-centre studies pooling magnetic resonance imaging (MRI) data across different scanners to advance the reproducibility of neuroscience research. In the present study, we developed an analysis strategy for <u>S</u>canner-<u>S</u>pecific <u>D</u>etection (SS-Detect) of SBPs in multi-scanner studies, and evaluated its performance relative to a conventional strategy.

**Methods:** In the first experiment, the SimTB toolbox was used to generate simulated datasets mimicking twenty different scanners with common and scanner-specific SBPs. In the second experiment, we generated one simulated SBP from empirical gray matter volume (GMV) datasets from two different scanners. Moreover, we applied two strategies to compare SBPs between schizophrenia patients’ and healthy controls’ GMV from two different scanners.

**Results:** The outputs of the conventional strategy were limited to whole-sample-level results across all scanners; the outputs of SS-Detect included whole-sample-level and scanner-specific results. In the first simulation experiment, SS-Detect successfully estimated all simulated SBPs, including the common and scanner-specific SBPs whereas the conventional strategy detected only some of the whole-sample SBPs. The second simulation experiment showed that both strategies could detect the simulated SBP. Quantitative evaluations of both experiments demonstrated greater accuracy of the SS-Detect in estimating spatial SBPs and subject-specific loading parameters. In the third experiment, SS-Detect detected more significant between-group SBPs, and these SBPs corresponded with the results from voxel-based morphometry analysis, suggesting that SS-Detect has higher sensitivity in detecting between-group differences.

**Conclusions:** SS-Detect outperformed the conventional strategy and can be considered advantageous when SBM is applied to a multi-scanner study.

## INTRODUCTION

Source-based morphometry (SBM) is a data-driven multivariate approach for identifying cross-subject covarying structural brain patterns (SBPs) and the subject-specific loading parameters of these patterns.^1,2^ While this approach was initially proposed as an extension and a multivariate alternative to voxel-based morphometry (VBM) of gray matter volume (GMV),^3^ it has also been implemented to construct SBPs based on cortical thickness,^4^ myelin volume fraction,^5^ and fractional anisotropy.^6^

To date, SBM has been predominantly used in single-scanner studies to investigate neuroanatomical differences between populations and neuroanatomical correlates of demographic or clinical characteristics.^7-10^ More recently, SBM has also been employed in collaborative studies,^11-14^ since pooling of multi-scanner magnetic resonance imaging (MRI) data from multiple sites has gained research momentum in the past decade.^15,16^ Theoretically, SBM assumes common SBPs and varying loading parameters across all subjects of the investigated cohort.^1,3^ Therefore, the conventional strategy to implement SBM in a multi-scanner setting is to directly concatenate data from the different scanners to form a single matrix as the input of independent component analysis (ICA);^3^ the outputs are whole-sample-level SBPs derived from the entire study sample across all scanners. The main limitations of this strategy is that it only yields whole-sample-level SBPs and it cannot ascertain scanner-specific variations. There is growing interest in studying individual variability in neuroscience,^17,18^ but the data-pooling nature of current SBM techniques precludes personalized SBPs detection.^2^ To align SBM with the goal of precision neuroscience, it is vital that we develop new ways to model scanner-specific SBPs, and infer more accurate estimates of subject-specific loading parameters. To address this challenge, we developed SS-Detect, a novel analysis strategy to detect scanner-specific structural brain patterns in multi-scanners studies, and used simulation experiments and real-world datasets to evaluate its performance against the conventional strategy.

## METHODS

### SBM analysis strategies for multi-scanner studies

Principal component analysis (PCA) and ICA are necessary for the SBM approach demonstrated in the present study so we begin with a brief description. PCA is usually used as a data reduction and de-noising method in ICA,^19^ it is typically carried out by computing the eigenvalue decomposition of the sample covariance matrix or by using singular value decomposition on the data. Since its introduction to functional MRI studies,^20^ ICA decomposition has been one of the primary approaches for analyzing brain imaging data. ICA is a statistical method used to discover hidden factors from a set of observed data such that the factors are maximally independent. In the context of its application on MRI data, ICA works by decomposing the mixed observed data (i.e., MRI data) into maximally spatially independent SBPs revealing patterns of variation that occur in MRI images.^3^ Each spatial SBP is associated with a loading parameter vector.

The conventional SBM analysis method for a multi-scanner study concatenates all the data from the different scanners to form a single matrix as the input for further analyses (Fig 1A), including PCA compression and ICA decomposition.^20^ It assumes that all scanner data have common spatial SBPs.^21^ In this case 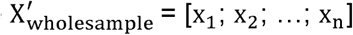 would be a typical data matrix, where **x**_***i***_ is the data matrix from the *i*-th scanner with dimension of *i*_*n*_×*v*, in which *i*_*n*_ is the *n*-th subject studied on the *i*-th scanner and *v* represents voxel number. In the present study, PCA via eigenvalue decomposition is performed on 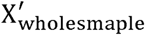 for dimensional reduction across all subjects scanned across all scanners. Thereafter, ICA is performed on X_wholesample_ which is the reduced data matrix, and the outputs of ICA are the whole-sample-level SBPs (S_wholesample_) and their associated loading parameter matrix (A_wholesample_): X_wholesample_ = A_wholesample_ S_wholesample_. Each column of the loading parameter matrix is a loading vector of the corresponding SBP presenting the weights of this SBP across all participants. More specifically, the loading parameter matrix expresses the relationship between all participants and SBPs: the columns of this matrix indicate how one SBP contributes to all participants, e.g., ICA decomposition of GMV data delineates structural SBPs based on the covariation of GMV among participants, and provides each participant an index (loading parameter) that reflects the degree to which each participant expresses the identified SBP. Unlike the VBM which assesses gray matter differences between individuals or groups by directly comparing their gray matter images, SBM assumes the consistency of the spatial structural brain patterns (SBPs) across the investigated participants, and the differences of the identified SBPs between different participants are expressed through the different loading parameters. In other words, one detects the gray matter differences of the SBPs by indirectly comparing loading parameters of different individuals or groups.

**Fig 1.**
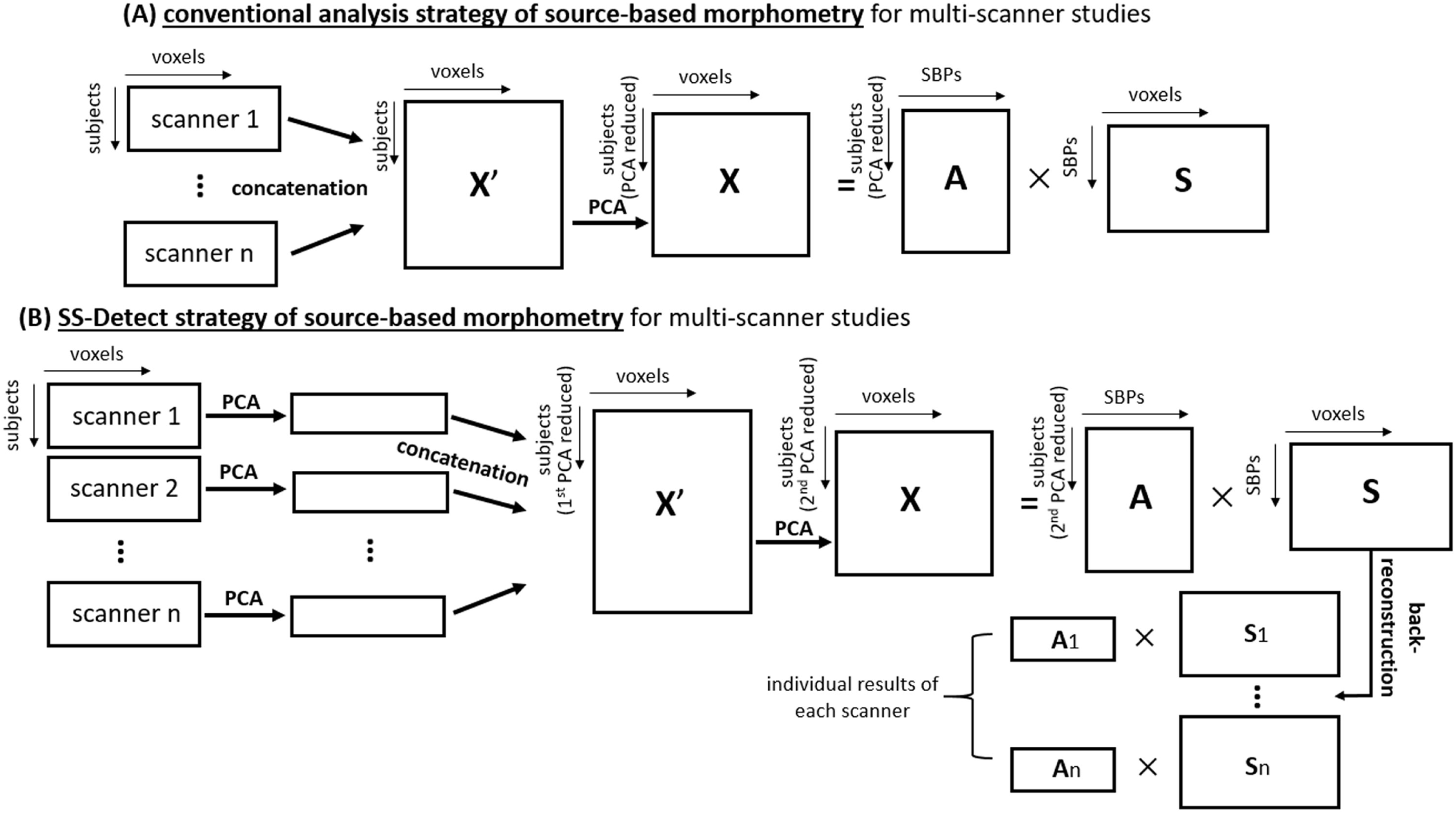
Flowcharts of the two analysis strategies of source-based morphometry (SBM). PCA: principal component analysis. X, A, and S indicate the MRI data matrix, loading parameter matrix, and spatial structural brain pattern (SBP) matrix. The MRI data matrix can be a matrix of gray matter volume, cortical thickness, or fractional anisotropy etc.

Fig 1B presents the flowchart of the SS-Detect which is a scanner-specific analysis strategy. By analogy to the group ICA approach used in functional MRI studies,^21^ SS-Detect first applies PCA to data matrix x_*i*_, *i* ∈ [1, n] at a scanner-level, then a second PCA procedure is conducted on the concatenated matrix 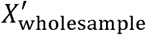 which is followed by the ICA decomposition procedure to obtain the whole-sample-level SBP matrix S_wholesample_ and loading parameter matrix A_wholesample_: X_wholesample_ = A_wholesample_ S_wholesample_, where X_wholesample_ is the output from the second PCA procedure and serves as the input of ICA decomposition. The final step is to back-reconstruct the scanner-specific-level results based on the whole-sample-level results.^22^ In the present study, dual regression was used to back-reconstruct scanner-specific SBPs and loading parameters. Given whole-sample-level S_wholesample_, dual regression first estimated scanner-specific loading parameter vectors, then scanner-specific SBPs, via multiple least-squares regression.^22,23^

### Simulation with SimTB toolbox

We evaluated and compared the conventional and the SS-Detect strategies with GMV data from simulated structural MRI data. GMV was used as an exemplar in the present study given that it is the most widely-used structural metric in the SBM literature,^1^ and GMV-based SBPs have been demonstrated to follow functionally meaningful architectures.^24,25^ However, the method can be implemented using other structural metrics (e.g. cortical thickness). We generated 20 GMV datasets mimicking 20 different scanners using the SimTB toolbox.^26,27^ In simulated data representing each scanner, the number of subjects was randomly generated between 50 and 100. Simulated data representing each scanner were generated as the product of a loading parameter matrix and a SBP matrix. The code used to generate simulated data is available on GitHub (https://github.com/ruiyangge/multiscanner_SBM). The first 10, and second 10 datasets were comprised of 15 SBPs each (with dimension 300×300 with baseline intensity of 1,000). The two sets of 10 datasets shared 14 common SBPs, yielding 16 different SBPs in total. The two dataset-specific SBPs were generated to mimic the variability of SBPs between different scanners. Spatial locations of these 16 SBPs were shown in Fig 2A. Rician noise was added to the simulated datasets with different signal-to-noise-ratio (SNR, uniformly varied from 40 to 110),^28,29^ defined as 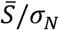, where 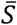 is the mean value of the SBP signal and σ_*N*_ is the standard deviation of the noise. Varying SNR values were used to mimic the varying distribution of SNR values across different scanners.^30^ Then, the conventional and the proposed analysis strategies were then applied to these 20 datasets, with model orders of ICA were set as 16 for both strategies. To ensure the stability of the ICA decomposition, the ICASSO technique with one-hundred ICA repetitions was used.^31^ Consistency of a spatial SBP estimated from different repetitions of ICA was quantified using the ICASSO cluster quality index *I*_*q*_ ranging from 0 to 1, with close-to-one *I*_*q*_ for a given SBP indicates that the SBP is consistent and stable across these repetitions.

**Fig 2.**
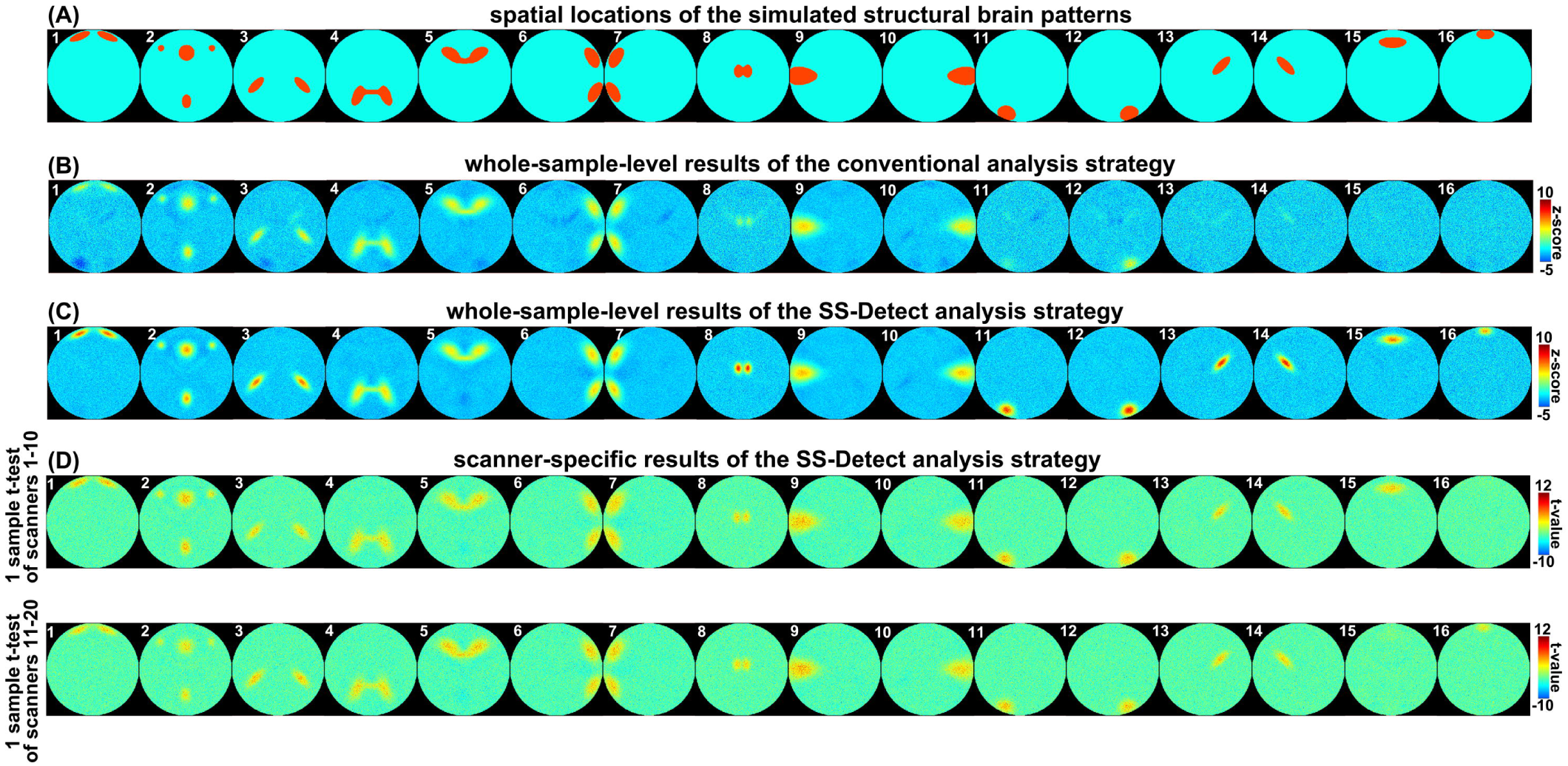
(A) Spatial locations of the simulated structural brain patterns; the first 10 data were comprised of 15 patterns (pattern IDs: 1-15), and the second 10 data were comprised of 15 patterns (pattern IDs: 1-14, and 16). Whole-sample-level results of the conventional analysis strategy (B) and SS-Detect analysis strategy (C). (D) Individualized scanner-specific results of the first 10 data and the second 10 data. A one-sample t-test was used to summarize the first 10 data and second 10 data separately.

### Simulation with empirical GMV data

In this simulation, we used two datasets collected at the University of British Columbia MRI Research Centre on a Philips Achieva 3.0-T scanner and a GE (General Electric) Genesis Signa 1.5-T scanner. Each dataset comprised 43 healthy participants (25 females). The age range of the participants was 18-63 and 16-47 years, for the Philips and GE datasets respectively. All participants provided written informed consent and studies were approved by the University of British Columbia ethics board. T1-weighted images from the Philips Achieva scanner were acquired with the following parameters: 165 axial slices; TR = 8.1 ms; TE = 3.5 ms; flip angle (FA) = 8°; field of view (FOV) = 256 mm × 256 mm × 165 mm; acquisition matrix = 256 × 250; slice thickness = 1 mm. T1-weighted images from the GE Genesis Signa scanner were acquired with the following parameters: 124 axial slices; TR = 11.2 ms; TE = 2.1 ms; FA= 20°; FOV = 256 mm× 256 mm × 260 mm; acquisition matrix = 256×256; slice thickness = 1.5 mm.

Preprocessing for VBM was performed using the CAT12 toolbox (http://www.neuro.uni-jena.de/cat/). First, the images were segmented into gray matter, white matter and cerebrospinal fluid probability maps by unified segmentation. Next, gray matter images were registered to the tissue probability map using affine transformation. Diffeomorphic Anatomical Registration using Exponential Lie Algebra was carried out to implement a high-dimensional nonlinear normalization. Through iteration of image registration and template creation, gray matter maps were normalized to their own average templates and further to the Montreal Neurological Institute space, and resampled to a voxel size of 1.5 mm^3^. Thereafter, normalized gray matter images were modulated with the Jacobian determinants of the nonlinear transformation and smoothed with an 8 mm^3^ Gaussian kernel. After preprocessing, we obtained normalized, modulated, and smoothed GMV images for subsequent SBM analysis.

We first decomposed each dataset into 10 SBPs with SBM, and replaced one randomly-selected empirical SBP with a simulated SBP located in the frontal lobe (Fig 3A). Specifically, data for the background voxels (voxels outside of the selected frontal region) were randomly generated with the MATLAB “randn” function, and data of the SBP voxels were drawn from the empirical SBP. The loading parameter matrix was randomly generated with the “randn” function. Simulated MRI data of each dataset were generated as the product of the loading parameter matrix and the SBP matrix. Thereafter, the conventional SBM analysis strategy, and the proposed strategy for multi-scanner studies were applied to the simulated datasets, with the model orders of ICA set as 10 for each strategy. This procedure was repeated ten times, with the ICASSO technique using one-hundred ICA repetitions in each run.

**Fig 3.**
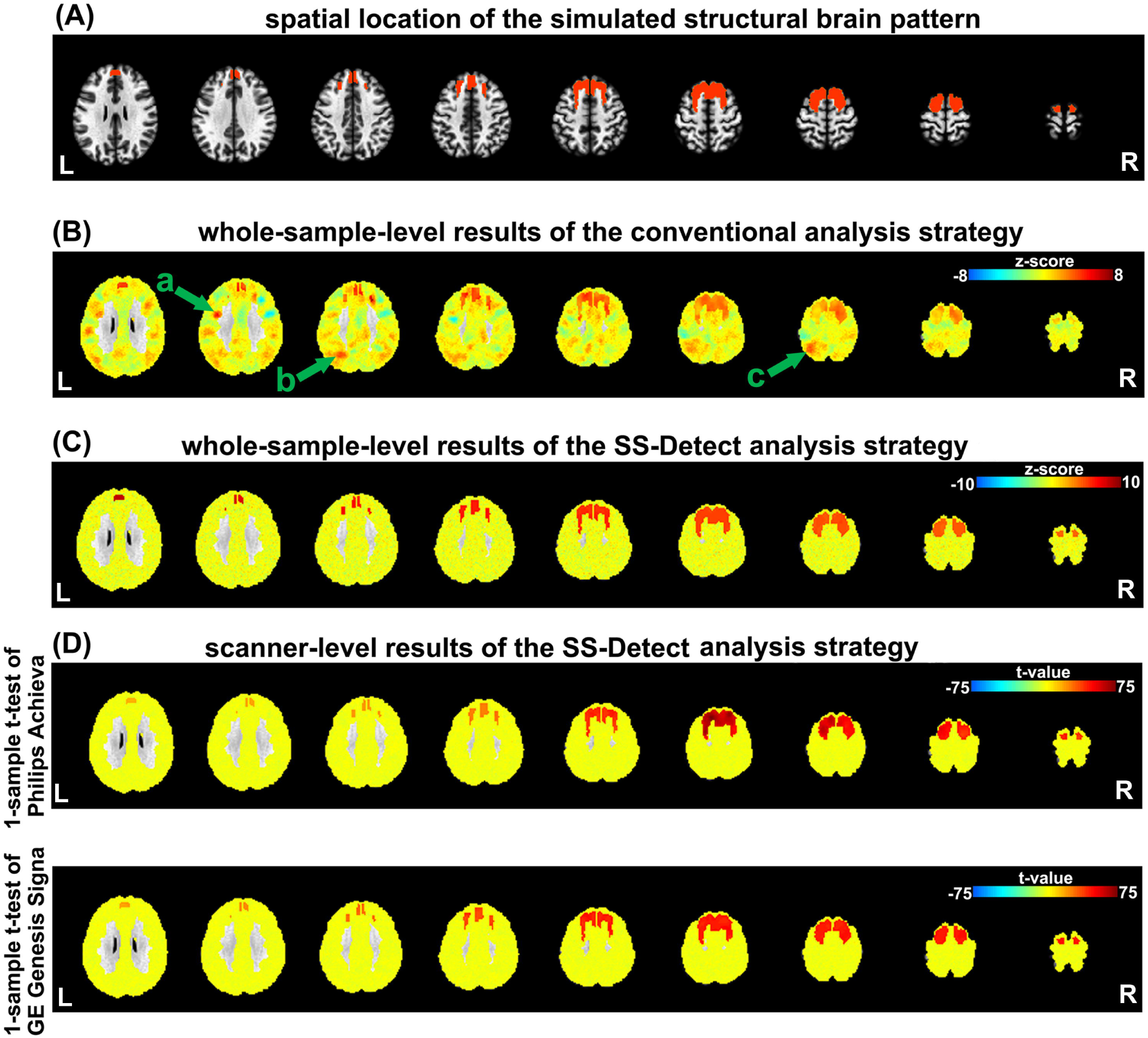
(A) Spatial location of the simulated component with the empirical gray matter volume data. Whole-sample-level results of the conventional analysis strategy (B) and SS-Detect analysis strategy (C). (D) Individualized scanner-specific results. A one-sample t-test was used to summarize the results of Philips and GE (General Electric) data separately. Green arrows in (B) indicate three large false-positive clusters detected by the conventional analysis strategy. L: left; R: right.

### Evaluation of the two strategies

Three quantitative measures were used to assess the two analysis strategies: (1) Dice’s coefficients between spatial templates and SBPs; (2) area under a curve (AUC) of the Dice’s coefficients curves; and (3) Pearson’s correlation coefficients between ground-truth loading vectors and estimated loadings.

In the SimTB-based simulated data, overall performance at the whole-sample-level results was the mean of each quantitative measure across the 16 SBPs; overall performance at the scanner-specific results was the mean of each quantitative measure across the 15 simulated SBPs for the first 10 and the second 10 datasets, respectively.

In the empirical data-based simulation, overall performance both at the whole-sample-level and the scanner-specific results was the final value of each performance measure averaged across ten runs.

Finally, we compared the performance measures (i.e., Dice’s coefficients, AUC of Dice’s coefficient’s curve, and correlation coefficients of loadings) between the two analysis strategies with Wilcoxon signed-rank tests. We used the ICASSO cluster quality index *I*_*q*_ to compare the consistency of the two strategies. Specifically, the mean *I*_*q*_ averaged over all SBPs was used for evaluation, and this was conducted on the whole-sample-level spatial SBPs on which the ICASSO repetitions were conducted. The statistical significance level was set as *p* < 0.05.

### Evaluation of the whole-sample-level spatial SBPs

The whole-sample-level spatial SBPs could be detected with both analysis strategies. For each experiment, we computed the Dice’s coefficients between the ground-truth templates (see Fig 2A, and Fig 3A) and the SBPs. Dice’s coefficient ranges from 0 to 1, with higher values indicating higher similarity between two binary images. We first z-transformed the SBPs, then used a z-threshold to binarize the SBPs, varying this threshold over an interval of [0.1, 10]. We took the Dice’s coefficient with z-threshold = 2.5 as one quantitative measure of the analysis strategies, this z-threshold was selected because it is a popular empirical threshold for displaying the results of spatial SBPs.^12,32-34^ We also plotted curves of the Dice’s coefficients with varying z-thresholds, and computed the area under curve (AUC) of the Dice’s coefficient curves as another quantitative measure.

### Evaluation of the scanner-specific spatial SBPs

The scanner-specific spatial SBPs could be detected only with SS-Detect. For the scanner-specific spatial SBPs of the SimTB data, we used a one-sample t-test to summarize the results of the first 10 and second 10 datasets separately, and used t-thresholds varying between 0.1 and 10 to binarize the t-maps.

For the scanner-specific spatial SBPs of the empirical data, we used a one-sample t-test to summarize the results of Philips and GE data separately, with t-thresholds varying between 0.1 and 75 to binarize the t-maps. Dice’s coefficient between the thresholded t-maps and ground-truth templates at each t-threshold was then computed.

### Evaluation of the loadings of the SBPs

Each subject had a single loading parameter for each SBP. To assess the detection ability of the analysis strategies for the loading parameters, we computed the Pearson’s correlations between the ground-truth loading vectors and those obtained from each strategy. This procedure was the same for both experiments.

### Application of the SS-Detect in a cross-sectional schizophrenia vs control study

The conventional and SS-Detect strategies were applied to GMV data of two schizophrenia datasets. One dataset (COBRE data; http://fcon_1000.projects.nitrc.org/indi/retro/cobre.html) was collected in a Siemens 3.0-T Trio TIM scanner with the following parameters: 192 sagittal slices; TR = 2,530 ms; TE = 1.64 ms; FA = 7°; FOV = 256 mm × 256 mm; acquisition matrix = 256 × 256; slice thickness = 1.0 mm. Given that there are differences in brain morphology between female schizophrenia patients and males schizophrenia patients,^35^ only MRI data from male participants were used in this experiment. This dataset consists of a group of 58 male schizophrenia patients (age MEAN±SD: 37.517±14.032) and a group of 51 male healthy controls (age MEAN±SD: 36.431±11.846). No significant difference was detected of the age (two-sample t-test, *p* = 0.666) between the two groups. The second dataset (UBC data) was collected in a GE Genesis Signa 1.5-T scanner with the following parameters: 124 axial slices; TR = 11.2 ms; TE = 2.1 ms; FA= 20°; FOV = 256 mm × 256 mm × 260 mm; acquisition matrix = 256 × 256; slice thickness = 1.5 mm. All participants in this dataset provided written informed consent and studies were approved by the University of British Columbia ethics board. This dataset consists of 48 male schizophrenia patients (age MEAN±SD: 20.908±4.060) and 39 healthy controls (age MEAN±SD: 23.559±10.168). No significant difference was detected of the age (two-sample t-test, *p* = 0.102) between the two groups. All schizophrenia patients in the two datasets were diagnosed on the basis of a structured clinical interview for DSM-IV-TR.

Preprocessing of voxel-based morphometry was performed using the CAT12 toolbox for each dataset separately, and each procedure was the same as the procedure mentioned in the simulation with empirical GMV data section. Subsequently, SBM analysis was conducted with conventional analysis strategy and SS-Detect strategy with the preprocessed GMV images as the inputs. The ICA order of each strategy was set to fifteen,^7^ and then by one-hundred ICASSO repetitions. We conducted cross-sectional comparisons between schizophrenia patients and healthy controls on the loading parameters of the SBPs through Wilcoxon rank-sum test, with age, total intracranial volume (TIV), and scanners as nuisance variables. Furthermore, we conducted between-group comparisons on the voxel-wise GMV images, with age, TIV, and scanners as nuisance variables. In these analyses, statistical significance level was set as *p* < 0.05.

## RESULTS

### Results from the simulated datasets

Fig 2 presents the estimated SBPs based on the two analysis strategies. At the whole-sample-level, the SS-Detect successfully detected all 16 simulated SBPs (Fig 2C), whereas the conventional analysis detected only 12 as some SBPs were apparently fused (Fig 2B); for example, one estimated SBP (pattern ID: 1, Fig 2B) consisted of two simulated SBPs (pattern IDs: 1 and 11, Fig 2A). SS-Detect successfully detected both the 14 common and the 2 scanner-specific SBPs in each scanner-specific dataset and at the whole-sample-level. The results of the first 10 datasets contained a “noise-like” SBP (pattern ID: 16) which was not simulated in these datasets, and the results of the second 10 datasets contained a “noise-like” SBP (pattern ID: 15) which was not simulated in these datasets.

Quantitative evaluation of the SBM results showed that at the whole-sample-level, estimated SBPs from the SS-Detect had higher AUC and Dice’s coefficient (z-score = 2.5) relative to the conventional analysis (Fig 4A). Although the first 10 simulated datasets had different SBPs compared with the second 10 datasets, their AUC and Dice’s coefficients were similar (Fig 4B). Fig 4C showed that the average correlation coefficient between the estimated loading parameters and the ground-truth loadings from SS-Detect was significantly higher than that from the conventional strategy. This finding was detected both in the first 10 datasets (MEAN±SD: 0.828±0.097 versus 0.745±0.317) and the second 10 datasets (MEAN±SD: 0.841±0.072 versus 0.731±0.344). The ICASSO cluster quality index *I*_*q*_ of SS-Detect was significantly higher than that of the conventional strategy (MEAN±SD: 0.856±0.171 vs 0.983±0.005, *p* = 0.006), demonstrating that SS-Detect produced more stable SBPs relative to the conventional strategy.

**Fig 4.**
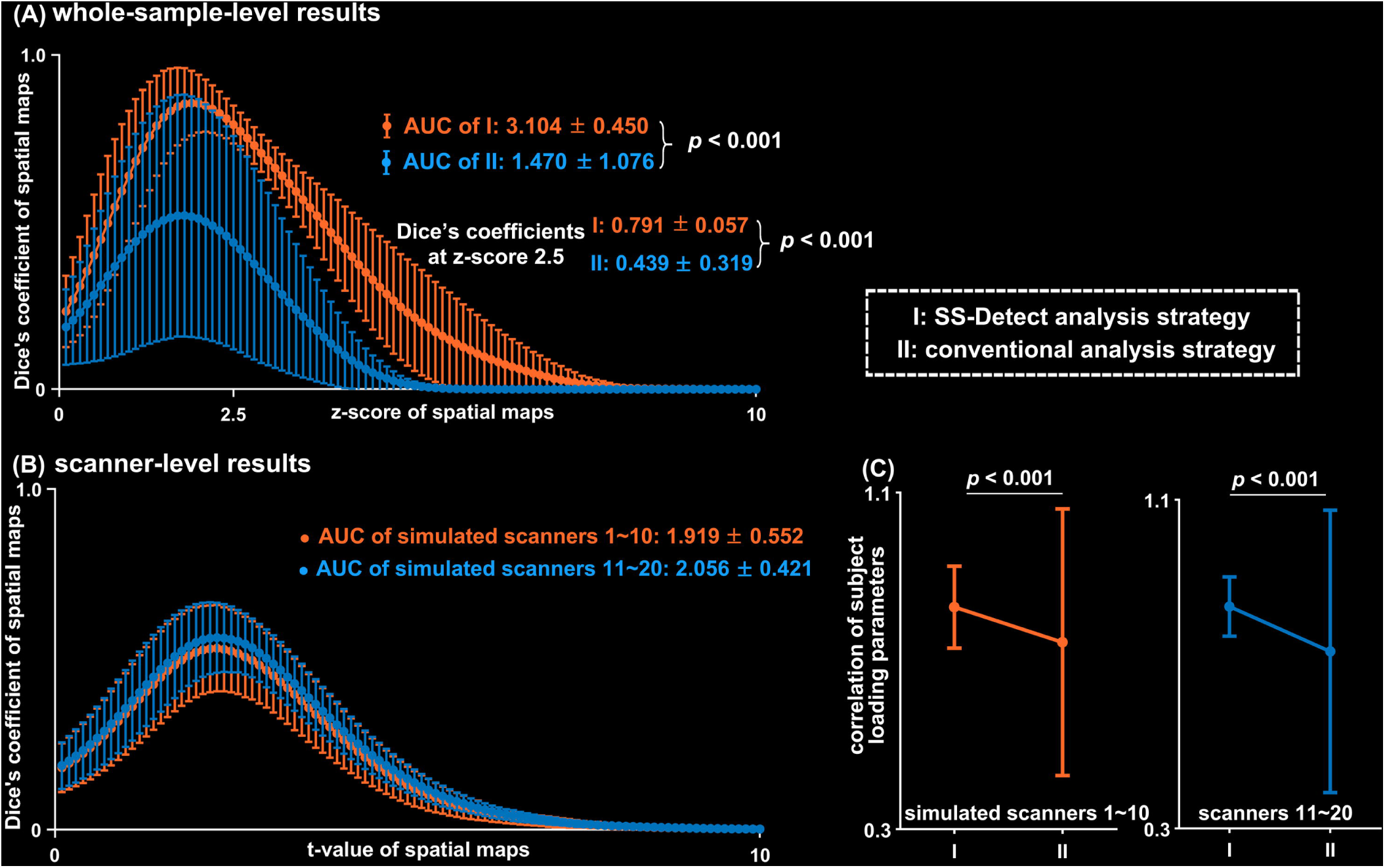
Results of the SimTB simulated datasets. (A) Average Dice’s coefficient between the 16 structural brain patterns (SBPs) and the ground truth at the whole-sample-level. Statistical analysis showed that the estimated SBPs from the SS-Detect analysis strategy had higher AUC and Dice’s coefficient (at z-score of 2.5) relative to the conventional analysis strategy. (B) Average Dice’s coefficient between the 15 SBPs and the spatial ground truth at the individual scanner-level for the first and second 10 simulated data, respectively. (C) Average correlation coefficient between the estimated loading parameters and the ground truth for the first and second 10 simulated data, respectively. Error bars indicate the standard deviation across all SBPs. AUC: area under curve.

### Results from the empirical data

For visualization purpose, we presented one exemplar result from the ten simulations of each analysis strategy at the whole-sample level. SS-Detect successfully estimated the simulated SBP (Fig 3C), whereas the estimated SBP from the conventional strategy included regions (e.g., left precentral lobe cluster ‘a’ and left superior parietal regions ‘b’ and ‘c’ in Fig 3B) which were not part of the SBP (Fig 3B). Therefore, the conventional strategy detected false-positive regions. We quantified the level of false-positive detection by counting the number of background voxels (voxels outside of the simulated frontal region) that were detected in the simulated SBP with threshold z > 2.5, i.e., if a background voxel had a z-value higher than 2.5, we counted it as a false-positive voxel. Results at the whole-sample-level spatial SBPs showed that average false-positive voxel count was 3320.100 (SD: 1782.621) and 15.700 (SD: 23.286) across the ten simulations for the conventional strategy and SS-Detect, respectively, and the statistical comparison between the two strategies on this count was significant (*p* < 0.001). At the scanner-level, the average false-positive voxel count across the ten simulations was 169.900 (SD: 165.721) and 129.200 (SD: 111.887) for the Philips Achieva scanner and GE Genesis Signa scanner, respectively. Quantitative evaluation of the SBM results showed that at the whole-sample-level, SS-Detect estimated SBPs with higher AUC and Dice’s coefficient (z-score = 2.5) relative to the conventional analysis (Fig 5A). At the scanner level, we presented the result from one-sample t-test of the ten simulations. SS-Detect successfully detected the simulated SBP for both datasets (Fig 3D). The AUC of the Philips dataset was slightly higher than that of the GE dataset (Fig 5B). Fig 5C showed that the average correlation coefficient between the estimated loading parameters and the ground-truth loadings from SS-Detect was significantly higher than that from the conventional strategy. This finding was observed in both in MRI data from Philips Achieva scanner (MEAN±SD: 0.877±0.070 versus 0.728±0.172) and GE Genesis Signa scanner (MEAN±SD: 0.895±0.055 versus 0.804±0.150). The ICASSO cluster quality index *I*_*q*_ of SS-Detect was significantly higher than that of conventional strategy (MEAN±SD: 0.972±0.052 vs 0.939 ±0.091, *p* = 0.002).

**Fig 5.**
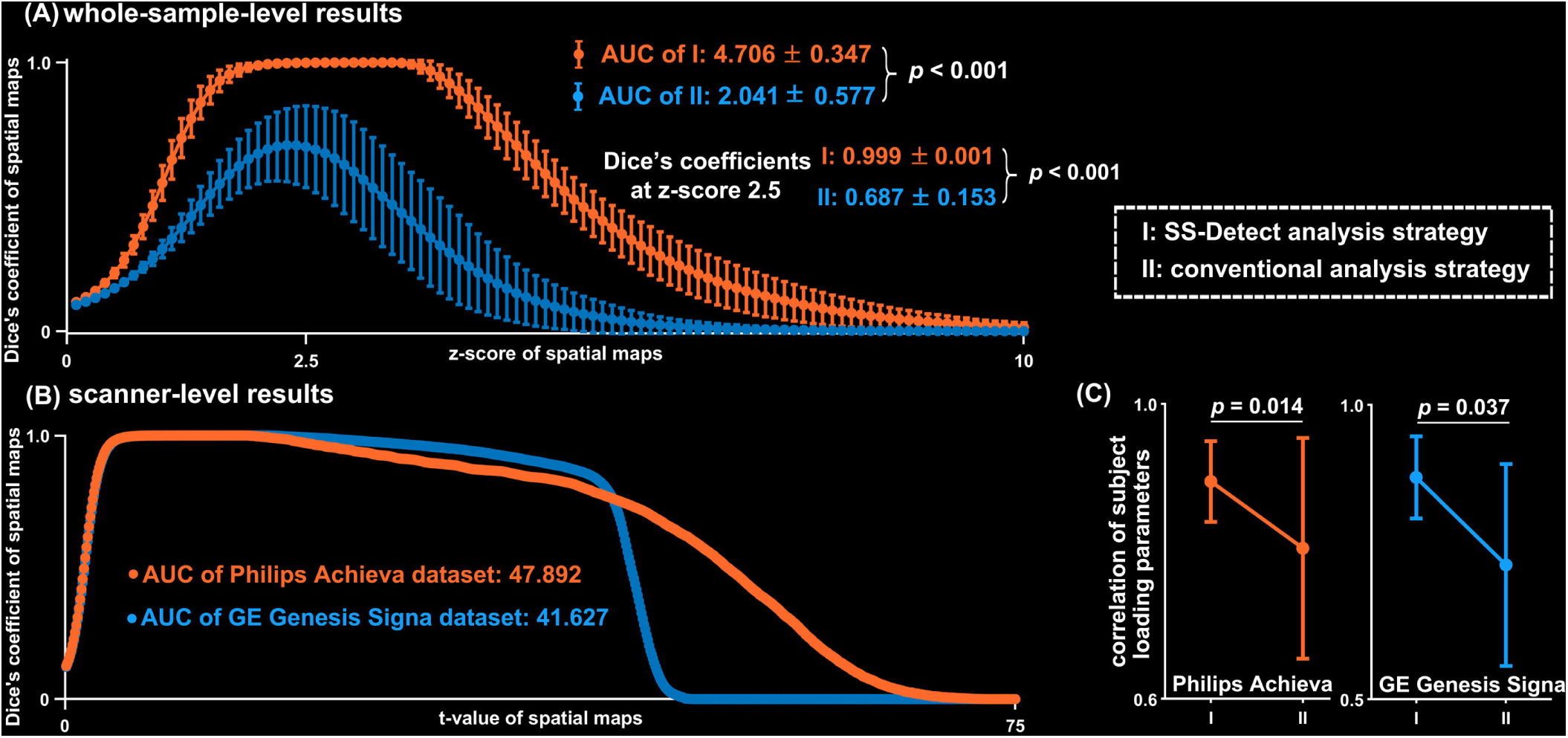
Results of the empirical datasets. (A) Average Dice’s coefficient between the simulated structural brain pattern (SBP) and the ground truth at the whole-sample-level. Statistical analysis showed that the estimated SBP from the SS-Detect analysis strategy had higher AUC and Dice’s coefficient (at z-score of 2.5) relative to the conventional analysis strategy. (B) Average Dice’s coefficient between the estimated SBP and the spatial ground truth at the individual scanner-level for the Philips and GE (General Electric) data, respectively. (C) Average correlation coefficient between the loading parameters and the ground truth for the Philips and GE data, respectively. Error bars indicate the standard deviation across the 10 repetitions of the simulation. AUC: area under curve.

### Results from the schizophrenia datasets

For each analysis strategy, spatial maps of fifteen SBPs were shown in Fig 6. Because ICA algorithms generate SBPs in an arbitrary order,^36^ and for visualization purpose, these SBPs were inspected and reordered based on the similarities between spatial maps from SS-Detect and conventional strategy. At the whole-sample-level, the outputs of the two SBM analysis strategies were highly similar except one cerebellar SBP (3^rd^ SBPs in Fig 6): SS-Detect strategy detected a SBP which located in the bilateral insula and anterior cingulate cortex; whereas the conventional strategy did not detect this SBP but one located in the precunes and posterior cingulate cortex, and this SBP largely overlapped with the 9^th^ SBP (Fig 6A). The ICASSO cluster quality index *I*_*q*_ of SS-Detect was higher than that of the conventional strategy (MEAN±SD: 0.980±0.006 vs 0.958 ±0.048, *p* = 0.051). Each SBP of the SS-Detect had an *I*_*q*_ higher than 0.950, suggesting the high reliability of these SBPs. For SBPs of the conventional strategy, thirteen had *I*_*q*_ higher than 0.950, and the 3^rd^ SBP had an *I*_*q*_ of 0.843 and 9^th^ SBP had an *I*_*q*_ of 0.840, suggesting that these two SBPs had lower reliability than other SBPs across the one-hundred ICASSO repetitions. All scanner-level SBPs from SS-Detect highly resembled to the corresponding whole-sample-level results, although the shape of the SBPs were slightly different between the two different datasets.

**Fig 6.**
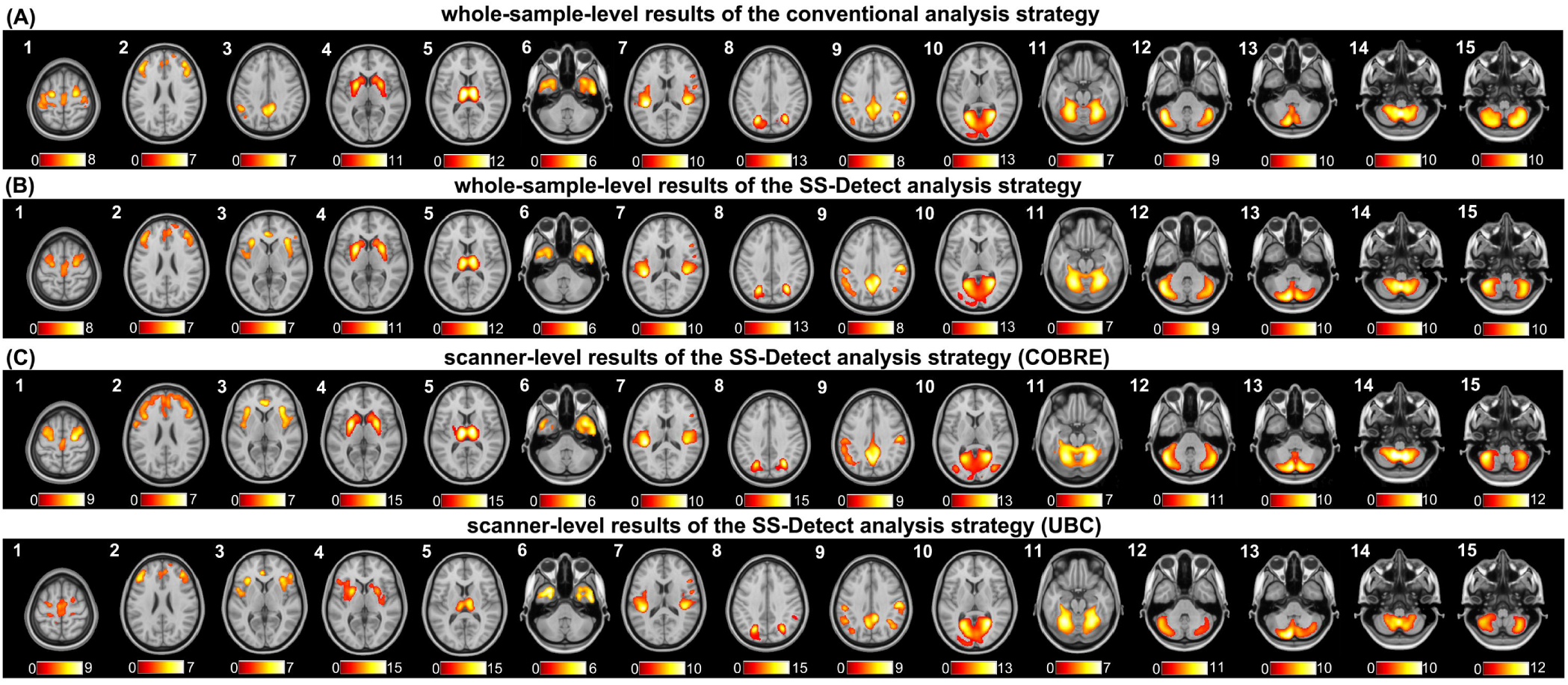
Spatial maps of the fifteen structural brain patterns (SBPs) projected over axial slices, at their respective global maximum coordinate. Spatial maps were thresholded at Z >□2.5 for visualization purposes, and a Z-score color bar was presented at the bottom of each SBP. L: left; R: right.

In the comparison between schizophrenia patients and healthy controls, both SS-Detect and the conventional strategy detected one SBP that controls showed significantly higher loading parameters than patients (Fig 7A and Fig 7B). This SBP located in bilateral superior temporal gyrus and posterior insula. SS-Detect detected three more SBPs that controls had significantly higher loading parameters: one SBP located in bilateral thalamus, one SBP located in bilateral middle prefrontal and medial prefrontal gyrus, and one located in bilateral superior, inferior and middle temporal gyrus, and temporal pole. The spatial location of these four SBPs overlapped with the VBM results (clusters ‘a’, ‘b’, ‘c’, and ‘d’ in Fig 7C). Fig 7D and 7E showed a cerebellar SBP that patients had significantly higher loading parameters than controls. This SBP located in cerebellar tonsil, and overlapped with the VBM results (cluster ‘e’ marked with green arrows in Fig 7F).

**Fig 7.**
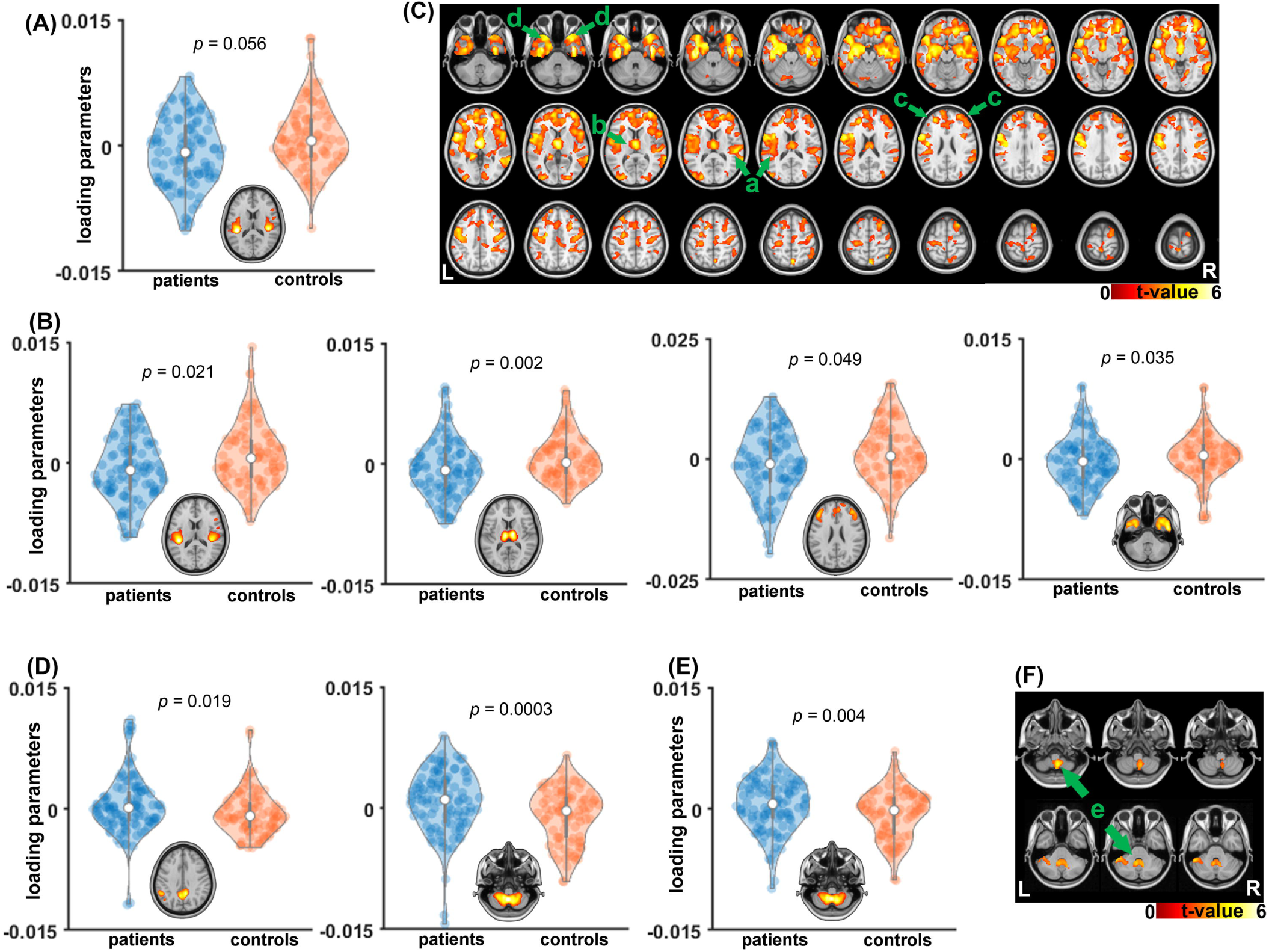
(A) One structural brain pattern (SBP) with significant higher loading parameters in the control group relative to the schizophrenia patient group was detected through the conventional analysis strategy. (B) Four SBPs with significant higher loading parameters in the control group relative to the schizophrenia patient group were detected through the SS-Detect strategy. *P*-value of between-group comparison for each SBP was shown. (C) Voxel-wise between-group comparison of the gray matter volume (GMV) images showed higher GMV in controls relative to patients (*p* < 0.05 uncorrected with cluster size *k* > 100). Four locations were marked with arrows, including [a] bilateral superior temporal gyrus and posterior insula; [b] bilateral thalamus; [c] bilateral middle and medial prefrontal gyrus; [d] bilateral superior, middle and inferior temporal gyrus, and temporal pole. (D) One precuneus/posterior-cingulate SBP and one cerebellar SBP with significant higher loading parameters in the schizophrenia patient group relative to the control group were detected through the conventional analysis strategy. *P*-value of between-group comparison for each SBP were shown. (E) One cerebellar SBP with significant higher loading parameters in the schizophrenia patient group relative to the control group was detected through SS-Detect. (F) Voxel-wise between-group comparison of the GMV images showed higher GMV in patients relative to controls (*p* < 0.05 uncorrected with cluster size *k* > 100). One cerebellar cluster [e] was marked with arrows. L: left; R: right.

## DISCUSSION

In this paper, we proposed SS-Detect as a scanner-specific analysis strategy to improve the detection of SBPs in SBM analyses of multi-scanner datasets. SS-Detect can be considered an adaptation to anatomical MRI studies of the commonly-used group ICA in the field of functional MRI.^21,37^ Both SS-Detect and the conventional strategy assessed in the present study employed group spatial ICA to establish correspondence of SBPs across datasets acquired with different MRI scanners. The present report describes a simulation study to assess the performance of SS-Detect. In the first simulation experiment, we demonstrated that SS-Detect could successfully estimate all SBPs, both at the whole-sample-level and at the individual scanner-level. These findings suggest that SS-Detect preserves the variability of the SBPs between different scanners, which cannot be ascertained with the conventional strategy. The conventional strategy did not recover all simulated SBPs correctly, even when some SBPs were shared between different datasets. This was because the conventional strategy concatenated the multi-scanner data along the subject dimension with an assumption that all scanner data have common spatial SBPs,^21^ and this assumption was inviolated by the fact that in our simulation we generated scanner-specific SBPs that were not shared by all scanners. Our results revealed that SS-Detect largely mitigated this constraint and successfully detected all SBPs. It is noted that for the SS-Detect results, Dice coefficients in the scanner-level were substantially lower than Dice’s coefficients in the whole-sample level. This phenomenon can be explained by the facts that with the current SS-Detect framework, scanner-level results were back-reconstructed based on the whole-sample-level results; in the meanwhile, the whole-sample-level results were produced by the concatenated data with larger sample size which could significantly increase statistical power and replicability relative to a single small-sample study.^38,39^ In the second simulation experiment, by using empirical GMV data, we demonstrated that SS-Detect detected less false-positive regions in the simulated SBP than the conventional strategy. Quantitative comparisons in two simulation experiments demonstrated that SS-Detect was more accurate than the conventional strategy in estimating both the spatial SBPs and subject-specific loading parameters. These findings underscore the advantage of using SS-Detect for analysis of SBM in a multi-scanner study.

Our experiment on schizophrenia patients and healthy controls showed that most of the SBPs generated by two different strategies were similar, and these SBPs have been revealed by previous studies with healthy participants.^25,40^ SS-Detect strategy detected a SBP which located in the bilateral anterior insula, and this SBP corresponded to the salience network.^41^ However, the conventional strategy did not detect this SBP but one (3^rd^ SBP in Fig 6A) largely overlapped with another SBP (9^th^ SBP in Fig 6A), this result signaled that the conventional strategy did not decompose the data in a proper way because spatial ICA used in the present study was meant to generate maximally independent SBPs, i.e., systematically spatially non-overlapping SBPs. Moreover, the lower reliability of these two SBPs across the ICASSO repetitions also suggested the deficiency of the conventional strategy relative to SS-Detect.

SBM was proposed as a multivariate alternative to voxel-wise VBM, and usually SBPs show significant between-group differences of SBPs that locate in brain regions that exhibit significant differences in the VBM analysis.^1,3^ In the comparison between patients and controls, both analysis strategies detected higher loading parameters of controls relative to patients in a superior temporal SBP, indicating GMV loss of this SBP in schizophrenia patients. This result has been consistently found in previous studies using SBM.^3,8,12^ More importantly, SS-Detect revealed three more SBPs (a thalamus SBP, a dorsolateral prefrontal SBP, and a temporal SBP in Fig 7B) that exhibited higher loading parameters in controls relative to patients, whereas the conventional strategy failed to detect these differences. Meta-analyses of the GMV studies suggested thalamic volume anomalies in schizophrenia patients,^42-44^ and SBM studies on schizophrenia patients also have reported decreased thalamic SBP in schizophrenia patients.^3,45^ Similarly, decreased GMV of bilateral middle prefrontal gyrus, inferior and middle temporal gyrus have been reported,^46-48^ and a multi-centre SBM study has discovered a middle prefrontal SBP decrement in schizophrenia patients.^12^ Our VBM analysis further confirmed schizophrenia patients’ GMV loss in the aforementioned regions (Fig 7C). All these results suggested that SS-Detect was more sensitive in detecting the GMV loss of SBPs than the conventional strategy. VBM analysis of the present study revealed a larger cerebellar tonsil GMV in patients relative to healthy controls (Fig 7F), and SBM results of both strategies confirmed this finding by showing that patients had higher loading parameters of the cerebellar tonsil SBP (Fig 7D and 7E). These results were contradictory to the results of a recent report in which cerebellar volume decrease was found in schizophrenia patients.^49^ Plausible explanations for this incongruence could be different gender distributions of the two studies,^50^ different medication status of the patients etc. Although it is beyond the scope of the present study which was to compare the two SBM analysis strategies, more work is needed to focus on the schizophrenia-related cerebellum abnormalities by disassociating different variables that might contribute to the inconsistency between different studies. Additionally, the conventional strategy detected a significant between-group difference in a precuneus/posterior cingulate SBP (Fig 7D), however, as mentioned earlier this SBP had low stability across the ICASSO repetitions and should be eliminated for further consideration as previous studies.^33^

Future studies will be necessary to extend this work in two ways. First, SS-Detect could be used to explore systematic differences introduced into a multi-scanner MRI study because of the different scanning platforms used; this could also assist in eliminating undesired noise which confounds true effects of interest.^51,52^ Second, relative to the conventional analysis strategy, SS-Detect is a further step towards the goal of precision neuroscience by establishing correspondence of SBPs in the entire study sample across scanners and simultaneously preserving the variability within each scanner-specific dataset. Nonetheless, the primary limitation of SS-Detect is that it cannot be applied at an individual subject level. Accordingly, future studies are need to go beyond, by addressing individual subject variability. Constructing SBPs from an individual anatomical MRI image would make it possible to investigate variability between individuals.^2,53^ Third, comparisons between schizophrenia patients and healthy controls were conducted on male participants only, therefore, findings in this experiment cannot be generalized to female population directly. Future studies are needed to characterize the gender-effects on the gray matter abnormalities in schizophrenia patients.

In conclusion, we proposed SS-Detect as a scanner-specific analysis strategy to improve the detection of SBPs in SBM analyses of multi-scanner datasets. With three experiments, we demonstrated that SS-Detect could successfully estimate all simulated SBPs, and detect less false-positive regions in a SBP, and improve the sensitivity in detecting between-group differences compared to the conventional strategy. Therefore, SS-Detect outperformed the conventional strategy and can be considered advantageous when SBM is applied to a multi-scanner study.

## Acknowledgements and Disclosure

We sincerely thank the study participants and MRI technologists at the UBC MRI Research Centre; we also sincerely thank the Center for Biomedical Research Excellence (COBRE) for sharing their data. RG, SD, TK, and SF declare no potential conflict of interest with respect to the research, authorship, and/or publication of this article. Dr. Honer has received consulting fees or sat on paid advisory boards for: AlphaSights, Guidepoint, In Silico, Translational Life Sciences, Otsuka, Lundbeck, and Newron, and holds shares in Translational Life Sciences and Eli Lilly. FVR reports research grants from CIHR, Brain Canada, Michael Smith Foundation for Health Research, and Vancouver Coastal Health Research Institute; reports receiving in-kind equipment support for this investigator-initiated trial from MagVenture; and consulting honoraria from Janssen pharmaceutical.

